# Benchmarking network propagation methods for disease gene identification

**DOI:** 10.1101/439620

**Authors:** Sergio Picart-Armada, Steven J. Barrett, David R. Willé, Alexandre Perera-Lluna, Alex Gutteridge, Benoit H. Dessailly

## Abstract

**Background:** In-silico identification of potential disease genes has become an essential aspect of drug target discovery. Recent studies suggest that one powerful way to identify successful targets is through the use of genetic and genomic information. Given a known disease gene, leveraging intermolecular connections via networks and pathways seems a natural way to identify other genes and proteins that are involved in similar biological processes, and that can therefore be analysed as additional targets.

**Results:** Here, we systematically tested the ability of 12 varied network-based algorithms to identify target genes and cross-validated these using gene-disease data from Open Targets on 22 common diseases. We considered two biological networks, six performance metrics and compared two types of input gene-disease association scores. We also compared several cross-validation schemes and showed that different choices had a remarkable impact on the performance estimates. When seeding biological networks with known drug targets, we found that machine learning and diffusion-based methods are able to find novel targets, showing around 2-4 true hits in the top 20 suggestions. Seeding the networks with genes associated to disease by genetics resulted in poorer performance, below 1 true hit on average. We also observed that the use of a larger network, although noisier, improved overall performance.

**Conclusions:** We conclude that machine learning and diffusion-based prioritisers are suited for drug discovery in practice and improve over simpler neighbour-voting methods. We also demonstrate the large effect of several factors on prediction performance, especially the validation strategy, input biological network, and definition of seed disease genes.

## Background

The pharmaceutical industry faces considerable challenges in the efficiency of commercial drug research and development [1] and in particular in improving its ability to identify future successful drug targets.

It has been suggested that using genetic association information is one of the best ways to identify such drug targets [2]. In recent years, a large number of highly powered GWAS studies have been published for numerous common traits (see for example [3] or [4]) and have yielded many candidate genes. Further potential targets can be identified by adding contextual data to the genetic associations, such as genes involved in similar biological processes [5, 6]. Biological networks and biological pathways can be used as a source of contextual data.

Biological networks are widely used in bioinformatics and can be constructed from multiple data sources, ranging from macromolecular interaction data collected from the literature [7] to correlation of expression in transcriptomics or proteomics samples of interest [8]. A large number of interaction network resources have been made available over the years, many of which are now in the public domain, combining thousands of interactions in a single location [9, 10]. They are based on three different fundamental types of data: (1) data-driven networks such as those built by WGCNA [8] for co-expression; (2) interactions extracted from the literature using a human curation process as exemplified by IntAct [11] or BioGRID [12]; and (3) interactions extracted from the literature using text mining approaches [13].

On the other hand, a plethora of network analysis algorithms are available for extracting useful information from such large biological networks in a variety of contexts. Algorithms range in complexity from simple first-neighbour approaches, where the direct neighbours of a gene of interest are assumed to be implicated in similar processes [14], to machine learning (ML) algorithms designed to learn from the features of the network to make more useful biological predictions (e.g. [15]).

One broad family of network analysis algorithms are the so-called Network Propagation approaches [16], used in contexts such as protein function prediction [17], disease gene identification [16] and cancer gene mutation identification [18]. In this paper, we perform a systematic review of the usefulness of network analysis methods for the purpose of identification of disease genes susceptible of being drug targets. Claims that such methods are helpful in that context have been made on numerous occasions but a comprehensive validation study is lacking. One major challenge in doing such a study is that it is not straightforward to define a list of known disease genes to be used for this purpose.

To address this, the Open Targets collaboration has been setup between pharmaceutical companies and public institutions to collect information on known drug targets and to help identify new ones [19]. A dedicated internet platform provides a free-to-use accessible resource summarising known data on gene-disease relationships from a number of data sources (e.g. known released drugs, genetic associations from GWAS, etc) [19].

The purpose of this work is to quantify the performance of network-based methods to prioritise novel targets, using various networks and validation schemes, and aiming at a faithful reflection of a realistic drug development scenario. We select a number of network approaches that are representative of several classes of algorithms, and test their ability to recover known disease genes by cross-validation.

We benchmark multiple definitions of disease genes, computational methods, biological networks, validation schemes and performance metrics. We account for all possible combinations of such factors and derive guidelines for future disease gene identification studies.

## Results

### Benchmark framework

Our general approach, summarised in figure 1, consisted in using a biological network and a list of genes with prior disease-association scores as input to a network propagation approach. We used three cross-validation schemes -two take into account protein complexes- in which some of the prior disease-association scores are hidden. The desired output was a new ranking of genes in terms of their association scores to the disease. Such ranking was compared to the known disease genes in the validation fold using several performance metrics. Given the amount of design factors and comparisons, the metrics were analysed through explorative additive models (see Methods). Alternatively, we provide plots on the raw metrics in the supplement, stratified by method in figures S10 and S11 or by disease in figures S12 and S13.

**Figure 1.**
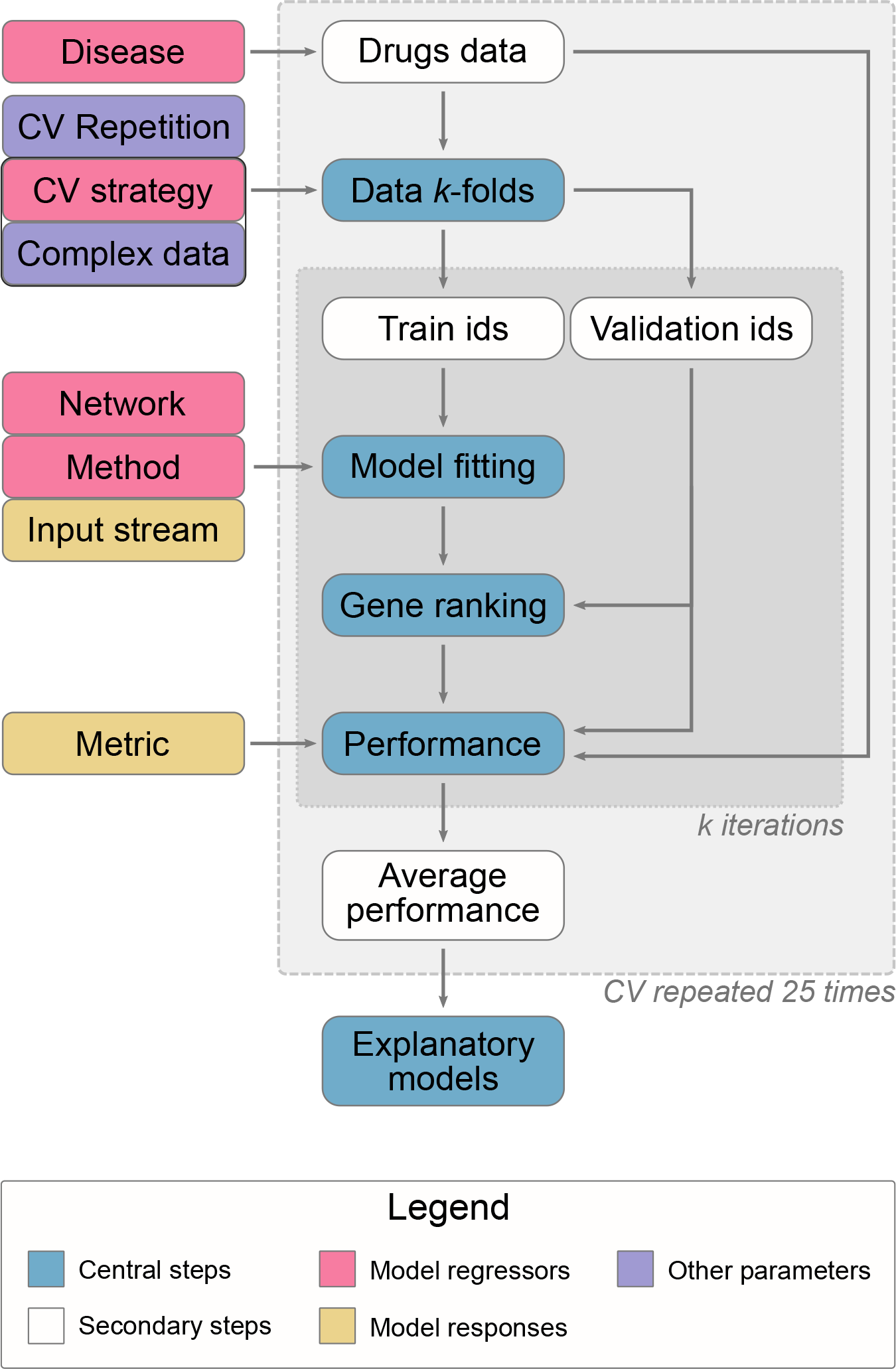
Benchmark overview. This work describes six performance metrics using two input streams (genetic association and drug-based genes) to predict drug-based genes for 22 common diseases. 3-fold cross validation (CV), repeated 25 times, was run under three CV strategies. The gene identifiers in each fold are determined using only the drugs data, regardless of the input. Two validation strategies are complex-aware and therefore needed this data to define the splits. 15 network-based methods (including 4 baselines) were evaluated, using two networks with different properties, by modelling their performance, averaged on every CV round. The explanatory models allowed hypothesis testing and a direct comparison between diseases, CV strategies, networks and methods.

We considered 2 metrics (AUROC and top 20 hits) and 2 input types (known drug target genes and genetically associated genes), resulting in a total of 4 combinations, each described through an additive main effect model. Another 4 metrics were explored and can be found in the supplement (figure S17 and tables S6, S7).

Interactions were explored, but they did not provide any added value for the extra complexity (see figure S18 from the supplement). The metrics used were the dependent variables, while the regressors included the prediction method, the CV scheme, the network and the disease.

### Performance using known drug targets as input

Figure 2 describes the additive models for AUROC and top 20 hits, and using known drug targets as input. Figure 3 contains their predictions for each method, network and cross validation scheme with 95% confidence intervals, averaged over diseases. The models are complex and we therefore review each main effect separately.

**Figure 2.**
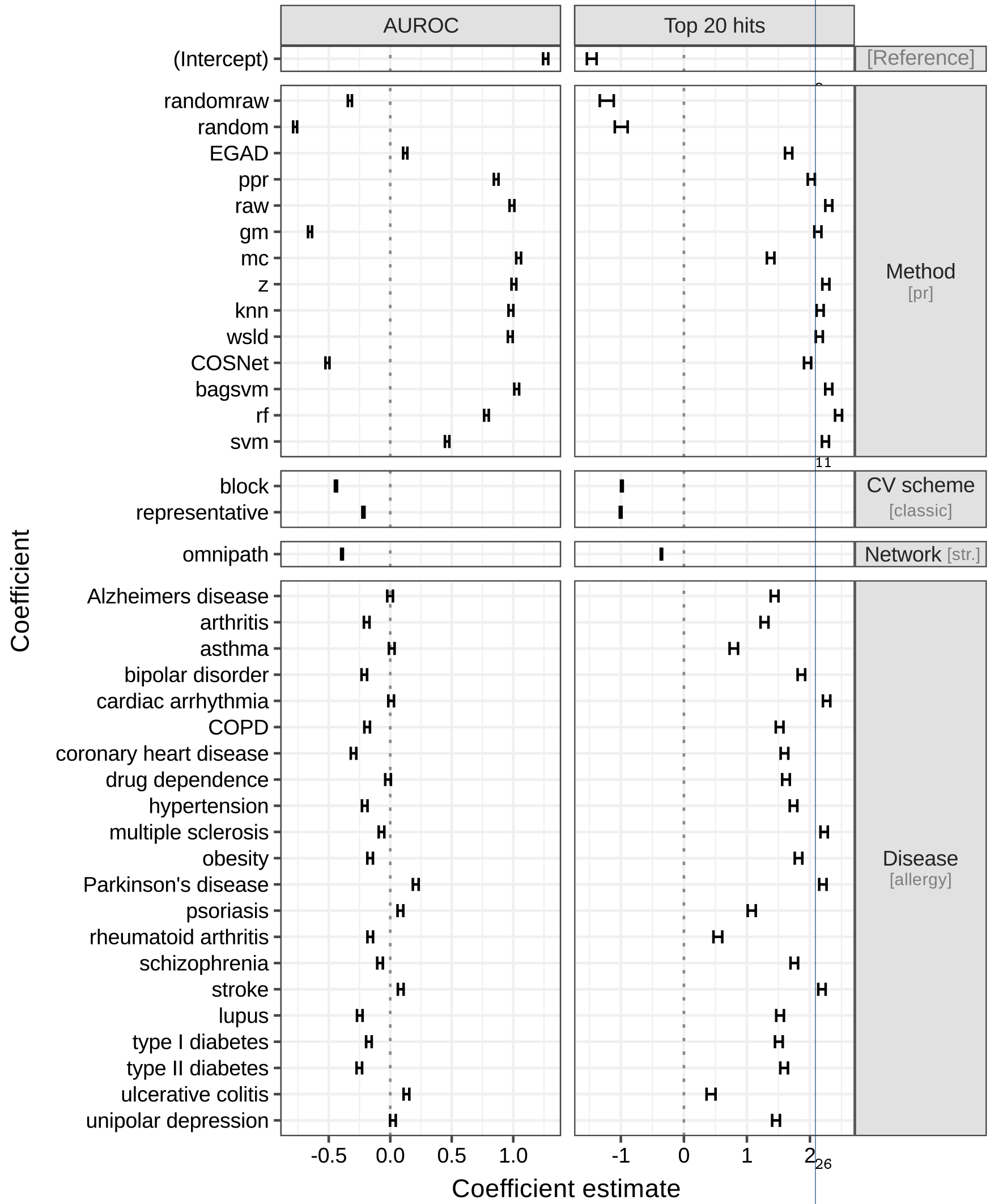
Additive models for AUROC and top 20 hits Each column corresponds to a different model, whereas each row depicts the 95% confidence interval for each model coefficient. Rows are grouped by the categorical variable they belong to: method, cv scheme, network and disease. Each variable has a **reference level**, implicit in the intercept and specified in brackets: pr method, **classic** validation scheme, **STRING** network and **allergy**. Positive estimates improve performance over the reference levels, whereas negative ones reduce it. For example, the data suggest that method rf performs better than the baseline using both metrics, and is the preferred method using the top 20 hits. Switching from STRING to the OmniPath network, or from classic to block or representative cross-validation, has a negative effect on both performance metrics. Specific model estimates and confidence intervals can be found in the supplement, see tables S8 and S9.

**Figure 3.**
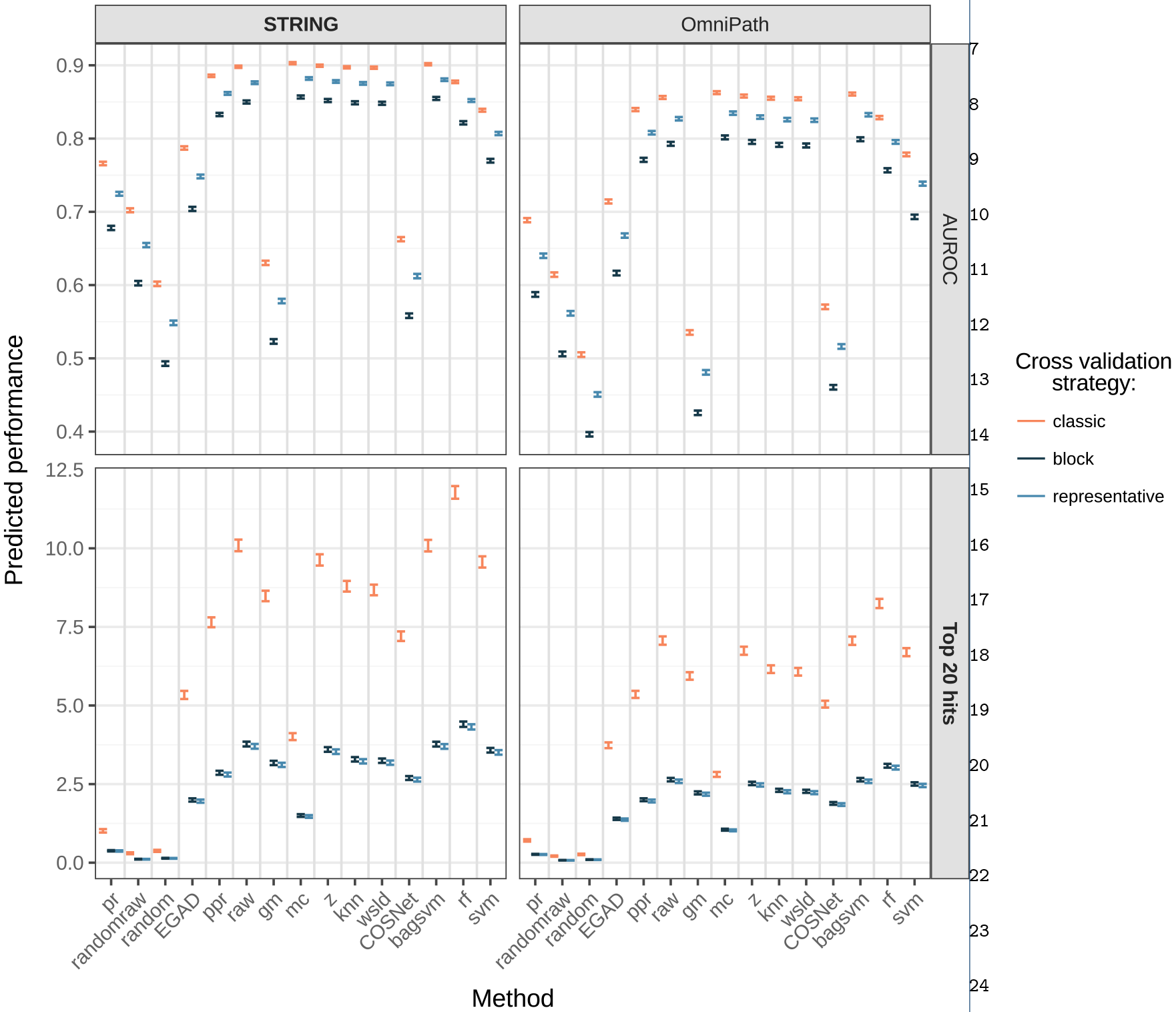
Performance predicted for AUROC and top 20 hits through the additive models Each row corresponds to a different model and error bars depicts the 95% confidence interval of the additive model prediction, averaging over diseases. In bold, the main network (STRING) and metrics (AUPRC, top 20 hits). A table with the exact values can be found in the supplement, table S9.

AUPRC (quasi-binomial model) and top 20 hits (quasi-poisson) behave alike, as can be observed by their similar ranking of model estimates in Figure 2. For inter-pretability within real scenarios, only top 20 hits is shown in the main body. The standard AUROC (quasi-binomial) clearly led to different conclusions and is kept throughout the results section for comparison. The remaining metrics (AUPRC, pAUROC 5%, pAUROC 10% and top 100 hits) result in similar method prioritisations as top 20 hits, see figure S17. Detailed models can be found in the supplement, indexed by tables S6 and S7.

#### Comparing cross-validation schemes

Whether protein complexes were properly taken into account when performing the cross-validation (see Methods) stood out as a key influence on the quality of predictions: there was a dramatic reduction in performance for most methods when using a complex-aware cross-validation strategy. For instance, method `rf` applied on the STRING network dropped from almost 12 correct hits in the top 20 predicted disease genes when using our *classic* cross-validation scheme down to fewer than 4.5 when using either of our complex-aware cross-validation schemes. Likewise, table S5 from the supplement ratifies that only the *classic* cross validation splits complexes. Our data suggests that the performance drop when choosing the appropriate validation strategy is comparable to the performance gap of competitive methods versus a simple neighbour-voting baseline (see figure 2). This highlights the importance of carefully controlling for this bias when estimating the performance of network-based disease gene prediction methods. Overall, the *classic* cross-validation scheme gave biased estimates in our dataset, whereas our *block* and *representative* cross-validation schemes had similar effects on the prediction performance. Method ranking was independent of the cross validation choice thanks to the use of an additive model. And since both the *block* and *representative* schemes make theoretical sense, we chose to focus on results from the block scheme in the rest of this study.

#### Comparing networks

We found that using STRING as opposite to OmniPath improved overall performance of network-based disease gene prediction methods. Our models for top 20 hits quantified this effect as noticeable although less important than that of the cross validation strategy. For reference, method `rf` obtains about 3 true hits under both complex-aware strategies in OmniPath. It has been previously shown that the positive effect on predictive power of having more interactions and coverage in a network can outweigh the negative effect of increased number of false positive interactions [20], which is in line with our findings. The authors also report STRING among the best resources to discover disease genes, which is a finding we reproduce here.

We focus on the STRING results in the rest of the text.

#### Comparing methods

Having identified the optimal cross-validation scheme and network for our benchmark in the previous sections, we quantitatively compared the performance of the different methods.

First, network topology alone had a slight predictive power, as method `pr` (PageR-ank approach that ignores the input gene scores) showed better performance than the `random` baseline under all the metrics. The randomised diffusion `randomraw` lied between `random` and `pr` in performance. Both facts support the existence of an inherent network topology-related bias among the positives that benefits diffusion-based methods.

Second, the basic GBA approach from `EGAD` had an advantage over the input-naïve baselines `pr`, `randomraw` and `random`. It also outperformed prioritizing genes using other Open Targets data stream scores such as genes associated to disease from pathways or from the literature (see supplement, table S19).

Most diffusion-based and ML-based methods outperformed `EGAD`. Results from top 20 hits suggest using `rf` for the best performance followed by, in order: `raw` and `bagsvm, z` and `svm` (main models panel in Figure 6).

To formally test the differences between methods, we carried a Tukey’s multiple comparison test on the model coefficients (Figure 5) as implemented in the R package multcomp [21]. Although such differences were in most cases statistically significant, even with such a strong multiplicity adjustment, their actual effect size or magnitude can be modest in practice, see Figures 3 and 6.

The ranking of methods was similar when using the metrics AUPRC, pAUROC and top *k* hits (see supplement, figure S17) and is only intended to be a general reference, given the impact of the problem definition, cross validation scheme and the network choice.

With AUROC on the other hand, `rf` lost its edge whilst most diffusion-based and ML-based methods appeared technically tied. Despite its theoretical basis, interpretability and widespread use in similar benchmarks, these results support the assertion that AUROC is a sub-optimal choice in drug discovery practical scenarios.

Figure 4 further shows how the different methods compare with one another. Distances between each pair of method in terms of their top 100 novel predictions were represented graphically. From this we observe that the supervised bagged SVM approach (`bagsvm`) behaves similarly to the simple diffusion approach (`raw`), reflecting the fact that they use the same kernel. We also observe that diffusion approaches do not necessarily produce similar results (compare for example `raw` and `z`). And that interestingly, methods `EGAD` (arguably one of the simplest) and `COSNet` (arguably one of the most complex) seemed to result in similar predictions. Fully supervised and semi-supervised approaches largely group in the top right hand quadrant of the STRING plot away from diffusion methods, possibly showing some shared greater potential for “learning effect” with the larger network.

**Figure 4.**
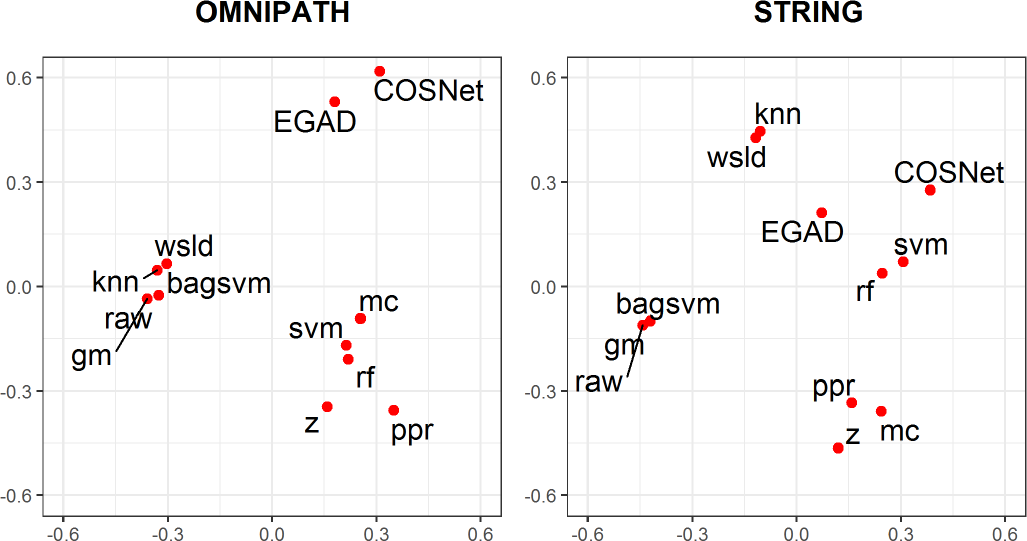
Multi-view MDS plot displaying the preserved Spearman’s footrule distances between methods. The differential ranking of their top 100 novel predictions using known drug target inputs are taken into account across all 22 diseases. Results are shown separately for the 2 networks considered in this study. Seed genes are excluded from the distance calculations.

**Figure 5.**
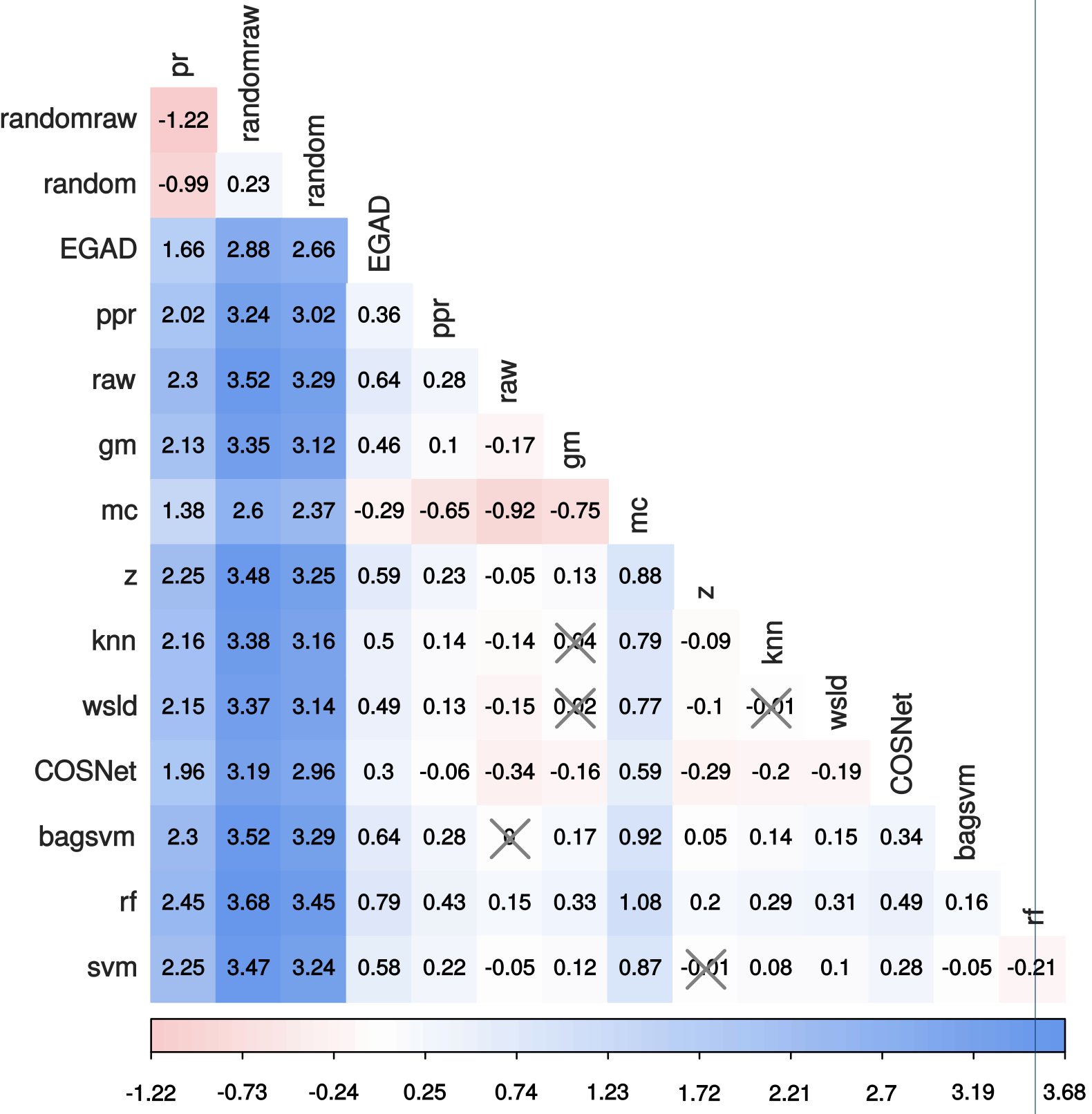
Pairwise contrasts on top 20 hits predicted by the quasipoisson model. Differences are expressed in the model space. Most of the pairwise differences are significant (Tukey’s test, p <0.05) – non-significant differences have been crossed out.

**Figure 6.**
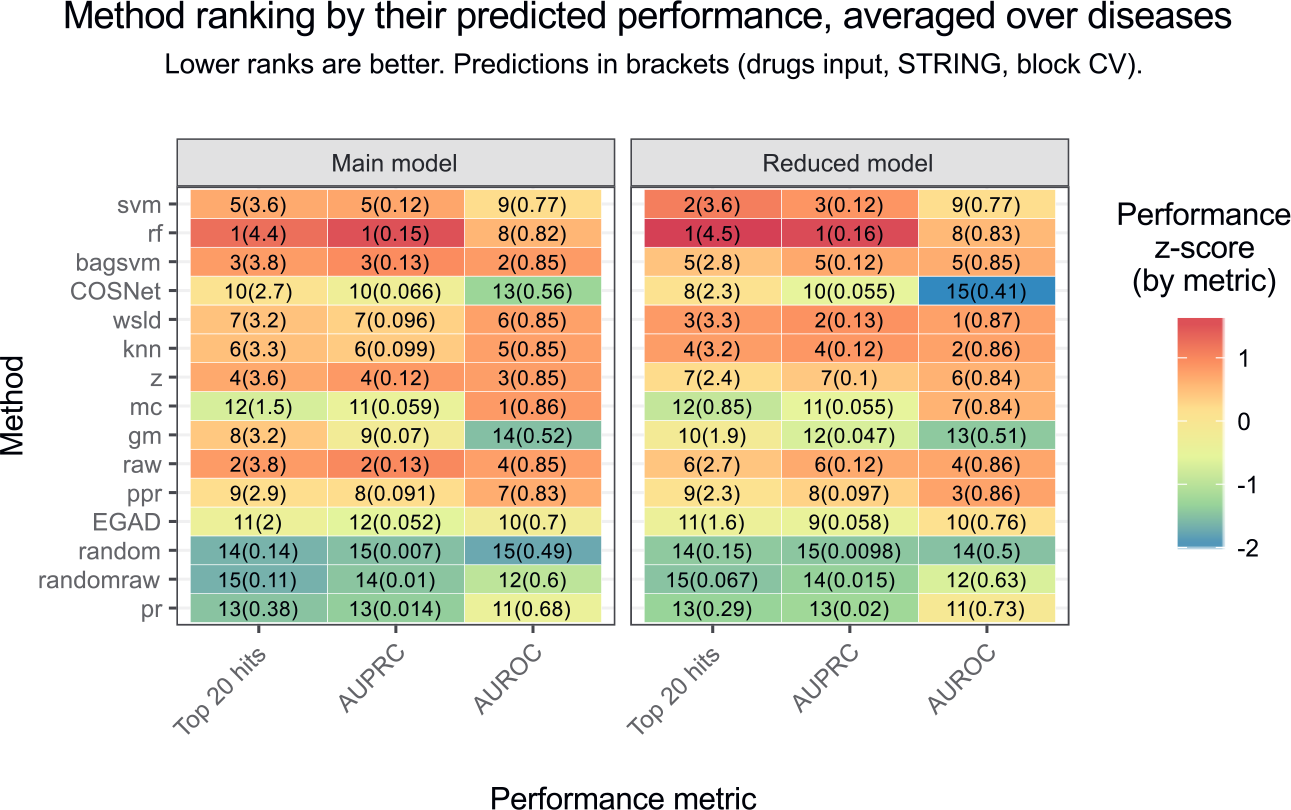
Ranking of all the methods, using the predictions of the main and the reduced models on the drugs input, STRING network, block cross validation and averaging over diseases. A column-wise z-score on the predicted mean is depicted, in order to illustrate the magnitude of the difference.

Interestingly, when comparing overall performances shown in figure 6 with the prediction differences from the MDS plot (figure 4), it appears that the better performing methods may be doing well for different reasons as they do not occur within the same region of the plot (e.g. `rf` and `raw`). MDS plots on the eight possible combinations of network, input type and inclusion of seed genes are displayed in the supplementary figures S15 and S16.

Regarding the STRING network and the block validation scheme, we fitted six additive models (one per metric) to the known drug target data (see supplement, table S7) and prioritised the methods (reduced models in figure 6). These reduced models better described this particular scenario, as they were not forced to fit the trends in all networks and validation schemes in an additive way. Considering the top 20 hits, `rf` and `svm` were the optimal choices, followed by `wsld` and `knn`.

#### Comparing diseases

We next examine performance by disease. The top 20 hits model in figure 2 shows that allergy (the figure’s baseline reference), ulcerative colitis and rheumatoid arthritis (group I) are the diseases for which prediction of disease genes was worst, whereas cardiac arrhythmia, Parkinson’s disease, stroke and multiple sclerosis (group II) are those for which it was best. As shown in figure 7, group I diseases had fewer known disease genes and lower modularity compared to group II diseases.

**Figure 7.**
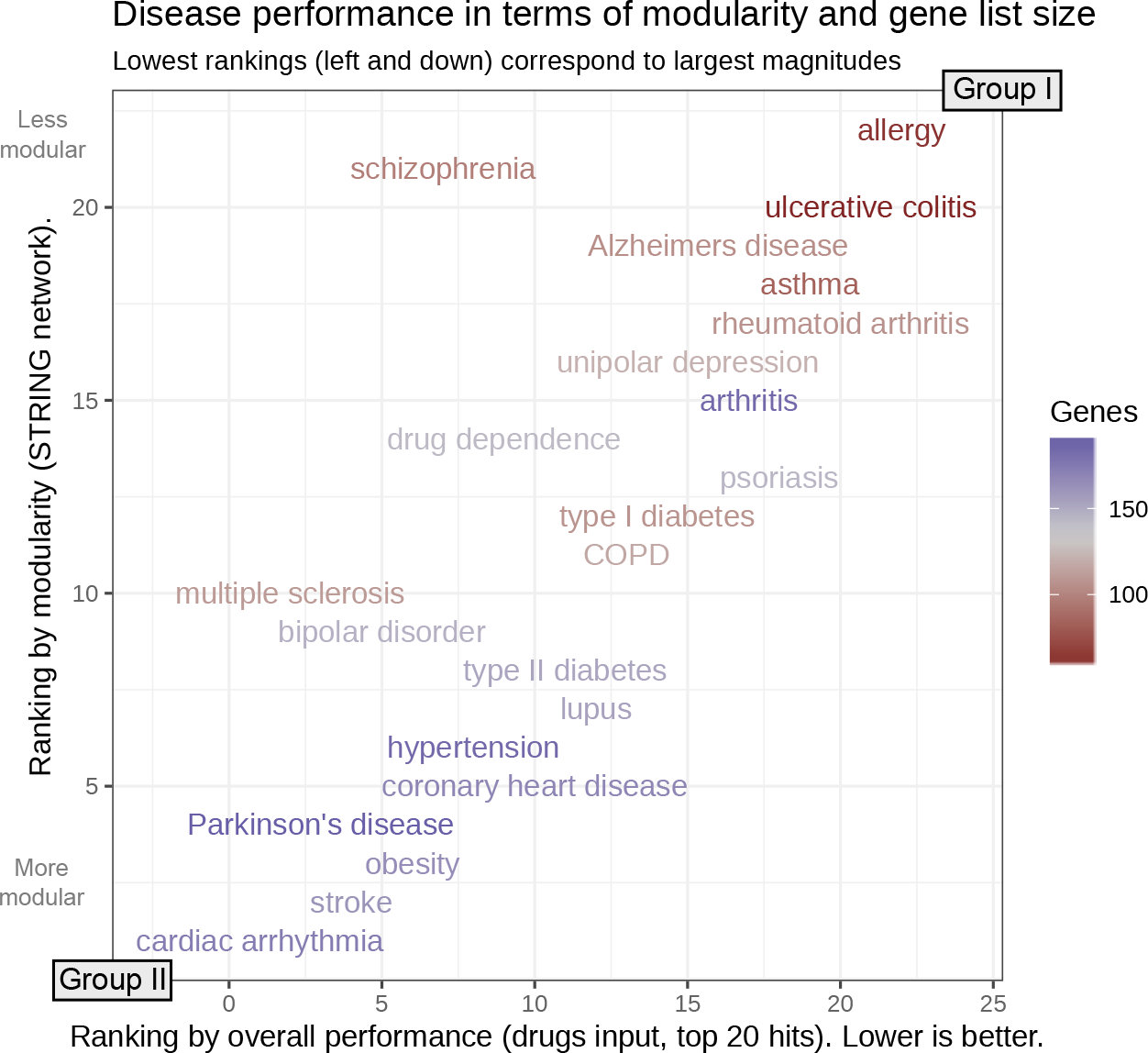
Disease performance ranked by the number of known disease genes from known drug data and modularity of known disease genes (obtained using the igraph package, see supplement, figure S6). Modularity is a measure of the tendency of known disease genes to form modules or clusters in the network. Diseases have been ranked using their coefficient from the top 20 hits metric with known drug targets as input (x axis) and their modularity (y axis). As discussed in the text, best predicted diseases tend to have longer gene lists and be highly modular.

Prediction methods worked better when more known disease genes were available as input in the network, with two possible underlying reasons being the greater data availability to train the methods, and the natural bias of top 20 hits towards datasets with more positives. Likewise, a stronger modularity within disease genes justifies the guilt-by-association principle and led to better performances. In turn, the number of genes and the modularity were positively correlated, see supplement, figure S14.

### Performance using genetic associations as input

Using genetically associated genes as input to a prediction approach to find known drug targets mimicked a realistic scenario where novel genetic associations are screened as potential targets. However, inferring known drug targets through the indirect genetic evidence posed problems to prediction strategies, especially those based on machine learning. Learning is done using one class of genes in order to predict genes that belong to another class, and the learning space suffers from intrinsic uncertainties in the genetic associations to disease.

Consequently, we observed a major performance drop on all the prioritisation methods: using any network and cross-validation scheme, the predicted top 20 hits were practically bounded by 1. This was more pronounced on supervised machine learning-focused strategies, as `rf` and `svm` lost their edge on diffusion-based strategies. The fact that the genetic associations of the validation fold were hidden further hindered the predictions and can be a cause of our pessimistic performance estimates.

#### Comparing cross-validation schemes

For reference, we also ran all three cross validation schemes on the genetic data to quantify and account for complex-related bias. The models confirm that, contrary to the drugs-related input, the differences between the results for the different cross-validation schemes were rather modest. For example, method `raw` with the STRING network attains 0.59-0.64, 0.50-0.54 and 0.37-0.40 hits in the top 20 under the classical, block and representative cross-validation strategies. The slightly larger negative effect on top 20 hits observed with the representative scheme is expected because the number of positives that act as validation decreased and this metric is biased by the class imbalance. The agreement between method ranking using AUPRC and top 20 hits was less consistent, possibly due to the performance drop, whilst AUROC again yielded quite a different ranking. Further data can be found in the supplement, tables S15 and S16.

#### Comparing networks

The change in performance for using the OmniPath network instead of the filtered STRING network was also limited. For AUROC the effect was negative, whereas for the top 20 hits metric the performance improved. Method `raw` changed from 0.50-top 20 hits in STRING to 0.61-0.66 in OmniPath under the block validation strategy.

#### Comparing methods

To be consistent with the drugs section, we take as reference the block cross-validation strategy and the STRING network.

The baseline approach `pr` that effectively makes use of the network topology alone proved difficult to improve upon, with 0.43-0.47 expected true hits in the top 20. Methods `raw` and `rf` respectively achieved 0.50-0.54 and 0.23-0.26 – although significant, the difference in practice would be minimal. The best performing method was mc with 0.65-0.7 hits. All the performance estimates can be found in the supplement, table S16. To give an idea of the effort that would be required in a realistic setting to find novel disease genes, the number of correct hits in the top 100 hits was 3.29-3.45 with the best performing method (in this case, `ppr`), against 2.25-2.38 of `pr`.

Two main conclusions can be drawn from these results. First, the network topology baseline retained some predictive power upon which most diffusion-based methods, as well as machine-learning approaches `COSNet` and `bagsvm`, only managed to add minor improvements, if any. Second, drug targets could still be found by combining network analysis and genes with genetic associations to disease, but with a substantially lower performance and with a marginal gain compared to a baseline approach that would only use the network topology to find targets (e.g. by screening the most connected genes in the network).

It is worth noting that gene-disease genetic association scores themselves have drawbacks and that better prediction accuracy could result as genetic association data improves.

## Conclusions

We performed an extensive analysis of the ability of network-based approaches to identify novel disease genes. We exhaustively explored the effect of different factors including the biological network, the definition of disease genes, and the statistical framework being used to evaluate methods performance. We show that carefully choosing an appropriate cross-validation framework and suitable performance metric have an important effect in evaluating the utility of these methods.

Our main conclusion is that network-based drug target discovery seems effective, reflecting the fact that drug targets tend to cluster within the network. This in turn may of course be due to the fact that the scientific community has so far been focusing on testing the same proven mechanisms. In a strict cross-validation setting, we found that even the most basic guilt-by-association method was useful, with ~2 correct hits in its top 20 predictions, compared to ~0.1 when using a random ranking. The best diffusion based algorithm improved that figure to ~3.75, and the best overall performing method was a random forest classifier on network-based features (~4.4 hits). Leading approaches can be notably different in terms of their top predictions, suggesting potential complementarity. We found a better performance when using a network with more coverage at the expense of more false positive interactions. In a more conservative network, random forest performance dropped to ~3.1 hits. Comparing performance on different diseases shows that the more known target genes, and the more clustered these are in the network, the better the performance of network-based approaches for finding novel targets for it.

We also explored the prediction of known drug target genes by seeding the network with an indirect data stream, in particular, genetic association data. Here, the best performing methods were diffusion-based and presented a statistically significant, but marginal, improvement over approaches that only look at network connectivity.

We conclude that network propagation methods can help identify novel disease genes, but that the choice of the input network and the seed scores on the genes needs careful consideration. Based on our approach and endorsed benchmarks, we recommend the use of methods employing representations of diffusion-based information (the MashUp network-based features and the diffusion kernels), namely random forest, the support vector machine variants, and raw diffusion algorithms for optimal results.

## Methods

### Selection of methods for investigation

Algorithms were selected for validation based on the following criteria:

1. published in a peer-reviewed journal, with evidence of improved performance in gene disease prediction relative to contenders,
2. Implemented As A Well Documented Open Source Package, That Is Efficient, RoBust And Usable Within A Batch Testing Framework,
3. directly applicable for gene disease identification from a single gene/protein interaction network, without requiring fundamental changes to the approach or additional annotation information and
4. capable of outputting a ranked list of individual genes (as opposed to gene modules for example).

In addition, we selected methods that were representative of a diverse panel of algorithms, including diffusion-based approaches, supervised learning approaches, and a number of baseline approaches (see table 1).

**Table 1.**
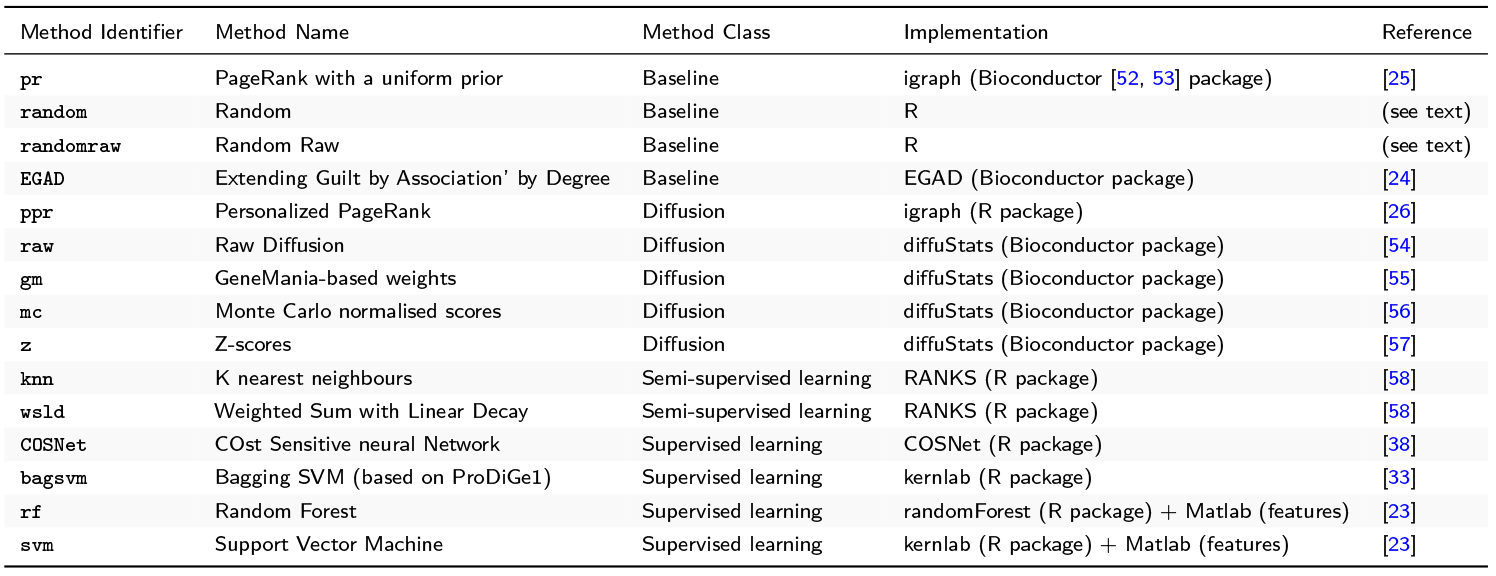
List of methods included in this benchmark. Method identifiers are shortened method names used throughout the text. Other columns are self-explanatory.

### Testing framework, algorithms and parameterisation

All tests and batch runs were set-up and conducted using the R statistical programming language [22]. When no R package was available, the methodology was re-implemented, building upon existing R packages whenever possible. Standard R machine learning libraries were used to train the support vector machine and random forest classifiers. Only the MashUp algorithm [23] required feature generation outside of the R environment, using the Matlab code from their publication. Further details on the methods implementation can be found in the supplement, section “Method details”.

EGAD [24], a pure neighbour-voting approach, was used here as a baseline comparator.

Diffusion (propagation) methods are central in this study. We used the random walk-based personalised PageRank [25], previously used in similar tasks [26], as implemented in igraph [27]. The remaining diffusion-based methods were run on top of the regularised Laplacian kernel [28], computed through diffuStats [29]. We included the classical diffusion `raw`, a weighted approach version `gm` and two statis-tically normalised scores (`mc` and `z`), as implemented in diffuStats. In the scope of positive-unlabelled learning [30, 31], we included the kernelised scores `knn` and the linear decayed `wsld` from RANKS [32]. Closing this category, we implemented the bagging Support Vector Machine approach from ProDiGe1 [33], here `bagsvm`.

Purer ML-based methods were also included. On one hand, network-based features were generated using MashUp [23] and two classical classifiers were fitted to them, based on caret [34] and mlr [35]. These are `svm`, the Support Vector Machine as implemented in kernlab [36], and `rf`, the Random Forest found in the randomForest package [37]. On the other hand, we tried the parametric Hopfield recurrent neural network classifier in the COSNet R package [38, 39].

Finally, we defined three naive baseline methods: (1) `pr`, a classic problem-naïve ‘non-personalised’ PageRank implementation where input scores on the genes are ignored; (2) `randomraw`, which applies the `raw` diffusion approach to randomly permuted input scores on the genes; and (3) `random`, a uniform re-ranking of input genes without any network propagation. The inclusion of `pr` and `randomraw` allowed us to quantify the predictive power of the network topology alone, without any consideration for the input scores on the genes.

### Biological networks

The biological network used in the validation is of critical importance as current network resources contain both false positive and false negative interactions, and these will affect any subsequent predictions [20].

Here, we used two human networks with different general properties, one more likely to contain false positive interactions (STRING [40]), and another more conservative (OmniPath [41]), to test the effect of the network itself on network propagation performance. We further filtered STRING [40] to retain only a subset of interactions. Having tested several filters, we settled upon high-confidence interactions (combined score > 700) with some evidence from the “Experiments” or “Databases” data sources (see supplement, table S2). Applying these filters and taking the largest connected component resulted in a connected network of 11,748 nodes and 236,963 edges. Edges were assigned weights between 0 and 1 by rescaling the STRING combined score.

We did not filter the OmniPath network [41]. After removing duplicated edges and taking the largest connected component, the OmniPath network contained 8,580 nodes and 42,145 unweighted edges.

### Disease gene data

We used the Open Targets platform [19] to select known disease-related genes. In this analysis we define disease-related genes are those reported in Open Targets as being the target of a known drug against the disease of interest, or as those with a genetic association of sufficient confidence with the disease. Associations were binarised: any non-zero drugs-related association was considered positive, implying that the methods would predict genes on which a drug has been essayed, regardless of whether the drug was eventually approved. Likewise, only genetic associations with an Open Targets score above 0.16 (see supplement, figure S1) were considered positive. We considered exclusively common diseases with at least 1,000 Open Targets associations, of which a minimum of 50 could be based on known released drugs and 50 on genetic associations, in order to avoid empty folds in the nested cross-validations. By applying these filters, we generated a list of phenotypes and diseases which we then manually curated to remove cancers (where causal genetic mechanisms can differ from those of other common diseases), non-disease phenotype terms (e.g. “body weight and measures”) as well as vague or broad terms (e.g.“cerebrovascular disorder” or “head disease”) and infectious diseases. This left 22 diseases considered in this study (table 2). Further descriptive material on the role of disease genes within the STRING network can be found in the section “Descriptive disease statistics in the STRING network” from the supplement.

**Table 2.** List of diseases included in this study. Diseases included in this study, with a minimum of 50 associated genes both in the known drug targets and the genetic categories (see text). The overlap between these two lists of genes showed a degree of dependence between these two Open Targets data streams for some of the diseases. P-values were calculated using Fisher’s exact test and are reported without and with correction for false discovery rate (Benjamini and Hochberg [59]).

| Disease | N(genetic) | N(drugs) | Overlap | P-value | FDR |
| --- | --- | --- | --- | --- | --- |
| allergy | 112 | 57 | 1 | 4.22e-01 | 4.42e-01 |
| Alzheimers disease | 208 | 103 | 4 | 1.10e-01 | 1.42e-01 |
| arthritis | 174 | 188 | 6 | 6.08e-02 | 1.03e-01 |
| asthma | 105 | 80 | 6 | 7.77e-05 | 5.70e-04 |
| bipolar disorder | 117 | 148 | 3 | 1.83e-01 | 2.12e-01 |
| cardiac arrhythmia | 75 | 177 | 6 | 9.15e-04 | 3.36e-03 |
| chronic obstructive pulmonary disease (COPD) | 154 | 116 | 6 | 4.18e-03 | 1.31e-02 |
| coronary heart disease | 111 | 171 | 4 | 7.86e-02 | 1.24e-01 |
| drug dependence | 75 | 143 | 6 | 2.96e-04 | 1.30e-03 |
| hypertension | 66 | 188 | 2 | 2.85e-01 | 3.14e-01 |
| multiple sclerosis | 71 | 167 | 4 | 1.83e-02 | 4.03e-02 |
| obesity | 69 | 194 | 3 | 1.06e-01 | 1.42e-01 |
| Parkinson's disease | 55 | 145 | 0 | 1 | 1 |
| psoriasis | 131 | 105 | 7 | 1.68e-04 | 9.23e-04 |
| rheumatoid arthritis | 138 | 95 | 5 | 5.18e-03 | 1.42e-02 |
| schizophrenia | 410 | 163 | 17 | 5.44e-05 | 5.70e-04 |
| stroke | 90 | 156 | 3 | 1.18e-01 | 1.44e-01 |
| systemic lupus erythematosus (lupus) | 126 | 109 | 5 | 6.30e-03 | 1.54e-02 |
| type I diabetes mellitus | 87 | 106 | 3 | 4.39e-02 | 8.04e-02 |
| type II diabetes mellitus | 130 | 154 | 4 | 9.14e-02 | 1.34e-01 |
| ulcerative colitis | 136 | 51 | 7 | 1.81e-06 | 3.98e-05 |
| unipolar depression | 123 | 121 | 4 | 3.81e-02 | 7.63e-02 |

### Validation strategies

#### Input Gene Scores

We used the binarised drug association scores and genetic association scores from Open Targets as input gene-level scores to seed the network propagation analyses (figure 8) and test their ability to recover known drug targets. With the first approach (subfigure (1) in figure 8), we tested the predictive power of current network propagation methods for drug target identification using a direct source of evidence (known drug targets). In the second approach (subfigure (2) in figure 8), we assessed the ability of a reasonable but indirect source of evidence – genetic associations to disease – in combination with network propagation to recover known drug targets.

**Figure 8.**
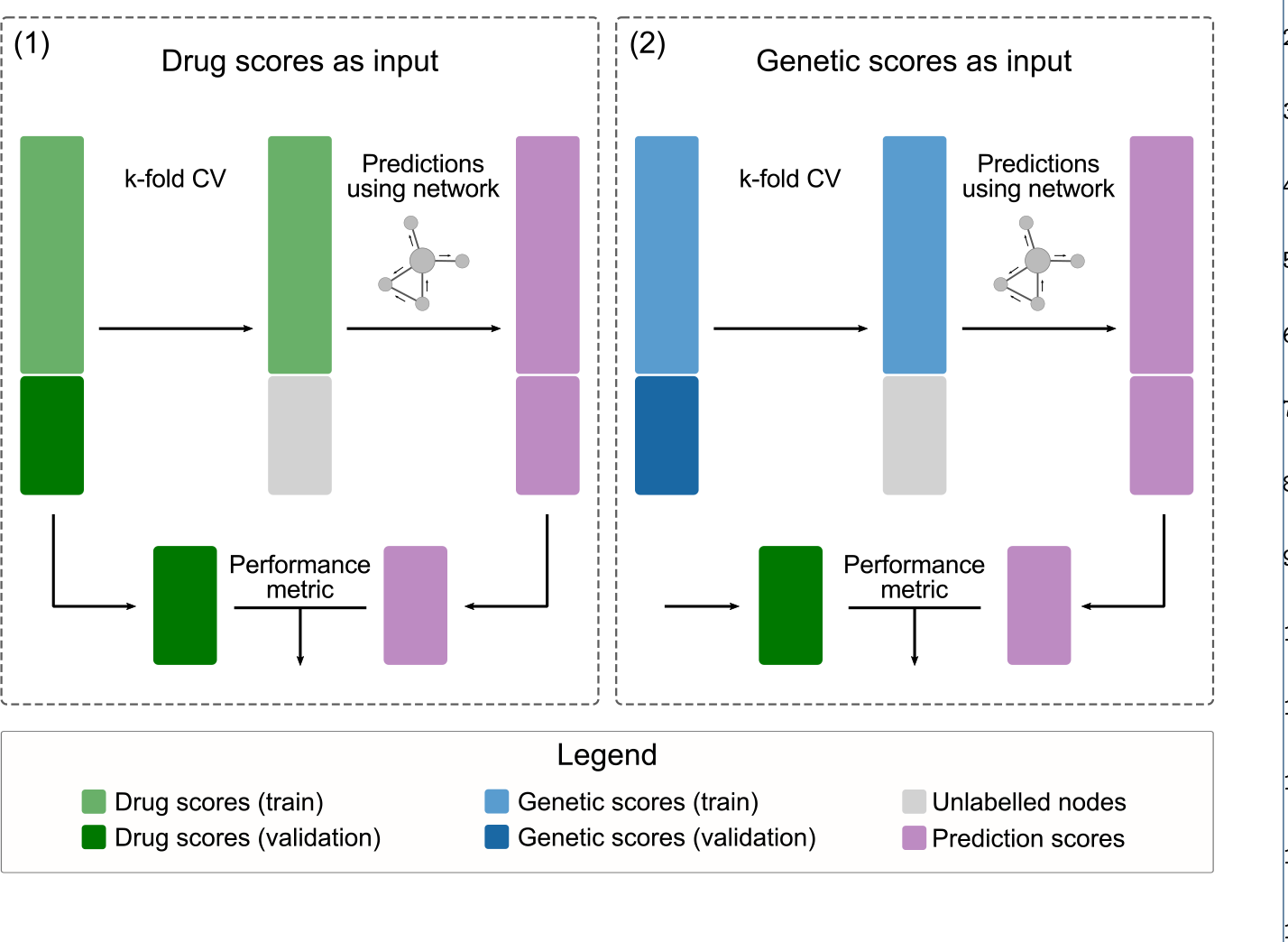
Input Gene Scores. Two input types were used to feed the prioritisation algorithms: the binary drug scores and the binary genetic scores. CV folds were always calculated taking into account the drugs input and reused on the genetic input.

#### Metrics

Methods were systematically compared using standard performance metrics. The Area under the Receiver Operating Characteristic curve (AUROC) is extensively used in the literature for binary classification of disease genes [42], but can be misleading in this context given the extent of the class imbalance between target and non-target genes [43]. We however included it in our benchmark for comparison with previous literature. More suitable measures of success in this case are Area under the Precision-Recall curve (AUPRC) [43] and partial AUROC (pAUROC) [44]. AUROC, AUPRC and pAUROC were computed with the precrec R package [45]. We also included top 20 hits, defined as the number of true positives in the top 20 predicted genes (proportional to precision at 20). It is straightforward, intuitive and most likely to be useful in practice, such as a screening experiment where only a small number of predicted hits can be assayed.

We considered another 3 metrics, reported only in Supplement, i.e. partial AUROC up to 5% FPR, partial AUROC up to 10% FPR, and number of hits within the top 100 genes.

#### Cross validation schemes and protein complexes

Standard (stratified) and modified k-fold cross-validation were used to estimate the performance of network-based methods. Folds were based upon known drugsrelated genes, regardless of which type of input was used (see figure 8). Genes in the training fold were negatively or positively labelled according to the input type, whereas genes in the validation fold were left unlabelled.

A fundamental challenge we faced when applying cross-validation to this problem was that known drug targets often consist in protein complexes, e.g. multiprotein receptors. Drug-target associations typically have complex-level resolution. The drug target data from Open Targets comes from ChEmbl [46], in which all the proteins in the targeted complex are labelled as targets.

If left uncorrected, this could bias our cross-validation results: networks densely connect proteins within a complex, random folds would frequently split positively labelled complexes between train and validation, and therefore network-based methods would have an unfair advantage at spotting positive hits in the training folds. In view of this, we benchmarked the methods under three cross validation strategies: a standard cross validation (A) in line with usual practice and two (B, C) complex-aware schemes (figure 9) addressing non-independence between folds when the known drug targets act as input.

**Figure 9.**
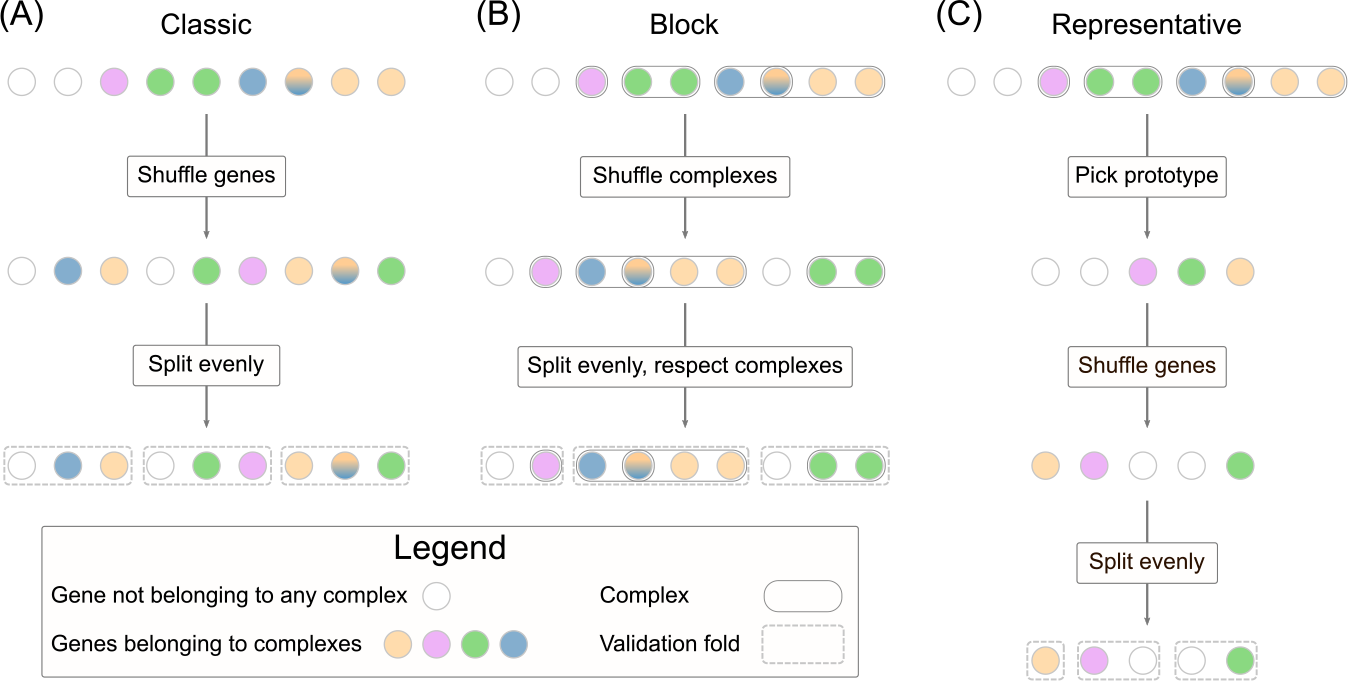
Cross-validation schemes. Three cross-validation schemes were tested. **(A)**: standard k-fold stratified cross-validation that ignored the complex structure. **(B)**: block k-fold cross-validation. Overlapping complexes were merged and the resulting complexes were shuffled. The folds were computed as evenly as possible without breaking any complex. **(C)**: representative k-fold cross validation. Overlapping complexes were merged and the resulting complexes from which unique representatives were chosen uniformly at random. Then a standard k-fold cross-validation was run on the representatives, but excluding the non-representatives from train and validation.

Strategy (A), called `classic`, was a regular stratified *k*-fold repeated cross-validation. We used *k* = 3 folds, averaging metrics over each set of folds, repeated 25 times (see also figure 1).

Strategy (B), named `block`, performed a repeated cross validation while explicitly preventing any complexes that contain disease genes to be split across folds. The key point is that, where involved, shuffling was performed at the complex level instead of the gene level – overlapping complexes that shared at least one known drug target were merged into a larger pseudo-complex before shuffling. Fold boundaries were chosen so that no complex was divided into two folds, while keeping them as close as possible to those that would give a balanced partition, see figure 9. Nevertheless, a limitation of this scheme is that it can fail to balance fold sizes in the presence of large complexes (see supplement, figure S9). For example, chronic obstructive pulmonary disease exhibited imbalanced folds, as 50 of the proteins involved belong to the Mitochondrial Complex I

Strategy (C), referred to as `representative`, selected only a single representative or prototype gene for each complex to ensure that gene information in a complex was not mixed between training and validation folds. In each repetition of cross validation, after merging the overlapping complexes, a single gene from each complex was chosen uniformly at random and kept as positive. The remaining genes from the complexes involved in the disease were set aside from the training and validation sets, in order (1) not to mislead methods into assuming their labels were negative in the training phase, and (2) not to overestimate (if set as positives) or penalise (if set as negatives) methods that ranked them highly, as they were expected to do so. This strategy kept the folds balanced, but at the expense of a possible loss of information by summarising each complex by a single gene at a time, reducing the number of positives for training and validation.

### Additive performance models

For a systematic comparison between diseases, methods, cross-validation schemes and input types, we fitted an additive regression model to the performance metrics of each (averaged) fold from the cross-validation. The use of main effect models eased the evaluation of each individual factor while correcting for the other covariates. We modelled each metric *f* separately for each input type, not to mix problems of different nature:

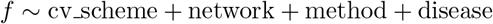

We fitted dispersion-adjusted logistic-like *quasibinomial* distributions for the metrics AUROC, pAUROC and AUPRC and *quasipoisson* for top *k* hits. *The effect of changing any of the four main effects is discussed in separate sub-sections in Results, following the order from the formula above*. After a data driven choice of cross-validation scheme and network we fitted reduced models within them for a more accurate description:

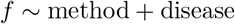

#### Qualitative methods comparison

The rankings produced by the different algorithms were qualitatively compared using Spearman’s footrule [47]. Distances were computed between all method ranking pairs for each individual combination of disease, input type, network and for the top *N* predicted genes, excluding the original seed genes. This part does not involve cross validation – all known disease-associated genes were used for gene prioritisations. Pairs of rankings could include genes uniquely ranked highly by a single algorithm from the comparison, so mismatch counts (i.e. percentage mismatches) between these rankings were also taken into account. Mismatches occur when a gene features in the top *N* predictions of one algorithm and is missing from the corresponding ranking by another algorithm. A compact visualisation of distance matrices was obtained using a multi-view extension of MDS [48, 49, 50]. For this we used the R package *multiview* [51] that generates a single, low-dimensional projection of combined inputs (disease, input and network).

## Competing interests

SB, DW, AG and BD are paid employees and shareholders of GlaxoSmithKline PLC. The commercial affiliation of SB, DW, AG and BD does not alter our adherence to BioMed Central policies.

## Availability of data and materials

The datasets supporting the conclusions of this article are available in https://github.com/b2slab/genedise.

## Author’s contributions

SP, SB, DW, and BD analysed and interpreted the data. AP, AG and BD helped supervise the project. All authors provided critical feedback and helped shape the research, analysis and manuscript. All authors approved the final version of this manuscript for publication.

## Acknowledgements

AG would like to acknowledge Philippe Sanseau, Matt Nelson and John Whittaker for critical feedback on the applicability of this research to drug discovery.

This work was supported by the Spanish Ministry of Economy and Competitiveness (MINECO) [TEC2014-60337-R and DPI2017-89827-R to AP].

AP and SP thank for funding the Spanish Networking Biomedical Research Centre in the subject area of Bioengineering, Biomaterials and Nanomedicine (CIBER-BBN), initiative of Instituto de Investigaciáon Carlos III 27(ISCIII). SP thanks the AGAUR FI-scholarship programme.

## Additional Files

Additional file 1 — Supplement

This document contains complementary material that supports our claims in the main body. It includes topics such as descriptive statistics, topological properties of disease genes, `raw` metrics plots, method details, MDS plots, alternative performance metrics and further explicative models.

Additional file 2 — MDS plots

Complementary single-disease MDS plots and distance matrices.

Additional file 3 — Interactions html viewer

Stand-alone viewer to explore models with interaction terms.

